# Nociception testing during fixed-wing ambulance flights. An interventional pilot study on the effects of flight-related environmental changes on the nociception of healthy volunteers

**DOI:** 10.1101/639781

**Authors:** Johannes Prottengeier, Stefan Elsner, Andreas Wehrfritz, Andreas Moritz, Joachim Schmidt, Michael Meyer

## Abstract

**Background:** The effects of environmental changes on the somato-sensory system during long-distance air ambulance flights need to be further investigated. Changes in nociceptive capacity are conceivable in light of previous studies performed under related environmental settings. We used standardized somato-sensory testing to investigate nociception in healthy volunteers during air-ambulance flights.

**Methods:** Twenty-five healthy individuals were submitted to a test compilation analogous to the quantitative sensory testing battery – performed during actual air-ambulance flights. Measurements were paired around the major changes of external factors during take-off/climb and descent/landing. Bland-Altman-Plots were calculated to identify possible systemic effects.

**Results:** Bland-Altman-analyses suggest that the thresholds of stimulus detection and pain as well as above-threshold pain along critical waypoints of travel are not subject to systemic effects but instead demonstrate random variations.

**Conclusions:** We provide a novel description of a real-life experimental setup and demonstrate the general feasibility of performing somato-sensory testing during ambulance flights. No systematic effects on the nociception of healthy individuals were apparent from our data. Our findings open up the possibility of future investigations into potential effects of ambulance flights on patients suffering acute or chronic pain.

## Introduction

Inter-hospital transfers are common medical procedures, that are sometimes carried out using fixed-wing air-ambulances. The number of such long-distance transfers is steadily rising due to the ongoing internationalization of specialized medical care and, much more importantly, due to increases in individual international mobility [1]. The latter results in growing numbers of aeromedical retrievals of travelers back to their home countries [2].

Long distance air ambulance flights can be considered a medical field of pre-requisites that truly distinguish it from intra-hospital care. While vibrations, noise, and restricted patient access must also be considered in other means of transportation, such as ground-ambulances and mobile ICUs, the rapid alterations in atmospheric pressure, oxygen partial pressure and air humidity that occur during airplane flights are environmental changes that are actually unique to this mode of transfer.

Despite this distinctiveness, most in-flight medical measures are simply extrapolated from what we know and do when on solid ground. For example, during transfers, analgesia is typically applied as if the patient were in a hospital – regardless of any of the possible effects, the profound environmental changes caused by flying in an airplane might have on human nociception.

Data from several studies have called this business-as-usual approach into question. For example, Sato and colleagues found that neuropathic pain was significantly aggravated in guinea pigs that were exposed to small alterations in atmospheric pressure similar to weather changes [3]. Additionally, healthy mountaineers in the Himalayas have been found to have lower pain detection thresholds when at high altitudes than when in low lying areas [4]. Thus, it seems that distinct environmental factors can influence nociception. And airplane travel in particular may affect other sensory functions as well. During simulated flights, healthy volunteers experienced changes in their gustatory detection thresholds [5]. As a consequence, commercial airlines have refined their in-flight meals to compensate for these flight-related sensory alterations.

In summary, it seems conceivable that airplane travel could impact nociception, but no data are available to evaluate its influence. In this prospective interventional study, we investigated the possible effects of air-ambulance flights on human nociception. Instead of artificially altering single environmental variables in a laboratory setting (such as atmospheric pressure), we decided to test pain perception in a real-life in-flight setting. This approach was used to provide external conditions identical to those encountered during medical transfers and to thus encompass the entirety of all possible influencing factors - even those, that can only be poorly simulated in a laboratory setting such as cabin noise, vibration etc..

## Materials and Methods

### Participants and setting

This study was approved of by the University of Erlangen-Nuremberg’s ethics council in advance under decision number 81_13 B.

The Department of Anesthesiology at Erlangen University Hospital is involved in international aeromedical retrievals as part of its cooperation with the ADAC, the German motorists club, which is one of the major insurance providers for Germans traveling abroad. The ADAC’s two Dornier 328 mid-range ambulance jets provided the setting for our experiments.

Healthy male volunteers were recruited from the pool of flight nurses and flight doctors engaged in transports on behalf of the ADAC. Informed written consent was obtained from each participant well before testing. All participants were required to undergo a concise health examination before they were included in our study. Exclusion criteria included, amongst others, any acute or chronic pain disorders, current or recent use of analgesics and any significant neurological, cardio-vascular, pulmonary or metabolic comorbidities.

### Test sequence

Nociception was tested at 4 distinct waypoints along the flight path. First, baseline values were obtained at ground level before take-off (Waypoint 1). Measurements for waypoint 2 were acquired after reaching cruising altitude. Waypoint 3 was set at a later time, right before leaving cruising altitude. Finally, a fourth and final set of measurements was obtained after touchdown, once the plane had reached its parking position (Waypoint 4). Picture 1 shows a schematic of the 4 waypoints along a flight.

Environmental factors such as atmospheric pressure, temperature and humidity were documented and were considered as possible influencing factors on nociception.

**SPACER Figure 1:**
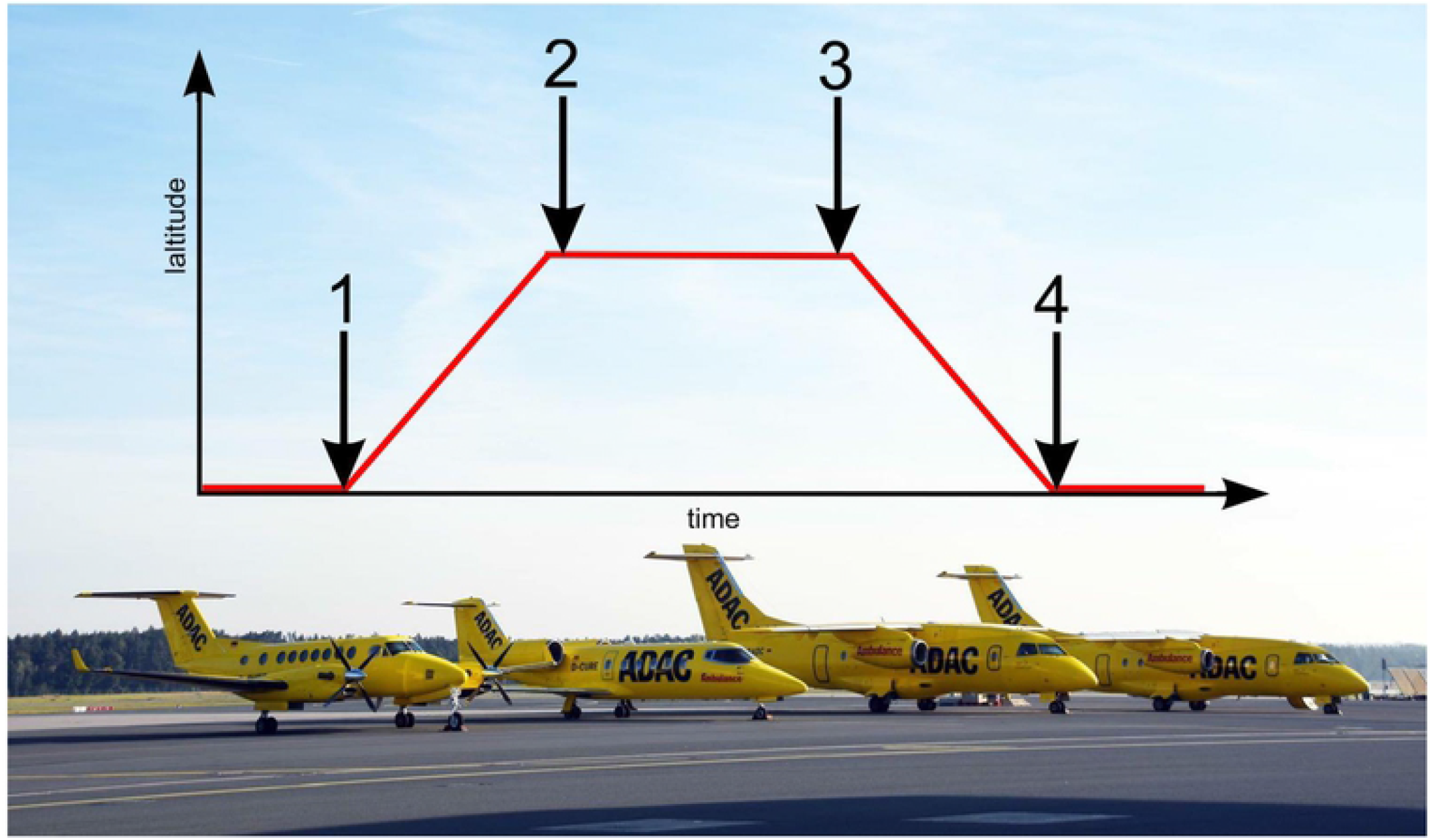
Schematic of test sequence. Legend Figure 1: The 4 sets of measurements were distributed strategically at distinct waypoints during each flight. Measurement 1 = before take-off, 2 = after reaching cruising altitude, 3 = before leaving cruising altitude, and 4 =after landing. The type of aircraft used for the experiments is shown in the background. The planes right and second to right are both identical Do 328s, in service as air ambulances [6].

### Quantitative sensory testing battery

Nociception measurements comprised a variety of modalities derived from the “quantitative sensory testing” (QST) battery. The QST has been developed by the German Research Network on Neuropathic Pain and has found widespread use worldwide since its introduction in 2002. Standardized testing allows the representative investigation of an individual’s somatosensory system, comprising both peripheral and central pathways [7–9]. The test procedures apply increasing, calibrated, non-invasive stimuli to detect the three distinct hallmarks of the sensory system for the different neurobiological sub-modalities of pain:

1. Perception thresholds,
2. Pain thresholds, and the
3. Quantification of sensations above threshold

Predefined techniques are provided by the QST manual to calculate validated threshold values from the obtained measurements. Briefly, QST measures through a set of tests (1.) when you first feel the stimulus, (2.) when the stimulus causes pain for the first time and (3.) how much a specific stimulus hurts.

### Thermal testing

Warm and cold thermal perception and pain thresholds were investigated using the TSA II NeuroSensory Analyzer (Medoc Advanced Medical Systems, Ramat Yishai, Israel). A thermode with a circulating water system was placed on the volunteer’s skin and a series of changes in water temperature were repeatedly applied. Technical limitations of thermode temperature were implemented to avoid skin lesions. When participants perceived that the temperature had changed and when they later felt pain derived from cold or heat, they pushed a button and the threshold temperatures were registered electronically.

### Mechanical testing

Mechanical pain thresholds were examined by means of pin-prick needle stimulators of increasing contact weights, resulting in stimulation intensities ranging from 8 Nm to 512 Nm against the intact skin surface of the participants. (Instruments were custom made by the expert mechanic workshop at the Department of Physiology, University of Erlangen-Nuremberg, Germany). Pain thresholds were derived from subjective oral ratings reported by the participants after repeated runs of stimulations. To detect the windup phenomenon, both single and series of above-pain-threshold stimuli were applied and rated on the numerical rating scale (NRS) for pain.

### Pressure algometer

The indenter-like pressure algometer FDN 200 (Wagner Instruments, Greenwich, USA) was pushed against the participant’s skin with increasing effort to determine the pressure pain threshold. The device was equipped with a pressure scale and readings were obtained when the volunteers verbally stated they perceived pain.

### Pain-Matcher

The pain matcher is not part of the QST test battery. It is a hand-held device that emits rectangular pulses of direct current between the participant’s first and second digitae [10, 11]. The transferred energy increases stepwise through automatic pulse elongation (60 steps from 0 to 450 msec.), resulting in an electrical sensation that becomes painful over time. The test subjects were instructed to loosen their grip on the device when thresholds were met. The intensity levels for perception and pain thresholds as well as for the individual’s maximum pain tolerance were displayed on the device and documented.

Table 1 provides a comprehensive overview of all test modalities. Abbreviations are later used in tables 3 and 4 of the results section.

**Table 1.** Synopsis of all test modalities and their abbreviations.

| **Test modality** | **Abbreviation** |
| --- | --- |
| Cold detection threshold | CDT |
| Heat detection threshold | HDT |
| Cold pain threshold | CPT |
| Heat pain threshold | HPT |
| Mechanical pain threshold | MPT |
| Wind up single stimulus | WUsS |
| Wind up multiple stimuli | WUmS |
| Pressure pain threshold | PPT |
| Pain matcher detection threshold | PMDT |
| Pain matcher pain threshold | PMPT |
| Pain matcher abort threshold | PMAT |

### Statistical analysis

To assess the influence of the environmental changes that occur between different flight phases on nociception, we performed Bland-Altman-analyses and prepared plots for every sensory test modality. Comparisons were paired around the phases of major changes in external conditions: take-off/climb and descent/landing. For this analysis, we matched waypoint 1 against waypoint 2 and waypoint 3 against waypoint 4. The solid black lines in the Bland-Altman-Plots represent the mean of the differences. The confidence intervals for means of differences are depicted as dashed black lines. The red upper (lower) lines show the upper (lower) limits of agreement equal to the mean ± 1.96SD. Usually, a total of 95% of observations lie within these limits. Confidence intervals for the limits of agreements were calculated and are presented in tables 3 and 4. However, for reasons of clarity, they were not included in the Bland-Altman-Plots. The blue lines represent a margin of ±20% around the means of the measurements obtained for each modality and serve as a possible indicator of clinical relevance.

Statistical analysis was performed using SPSS Statistics 21 (IBM Corp. Armonk, NY, USA). Values are presented as means with standard deviations and 95% confidence intervals, where appropriate.

## Results

### Descriptive statistics

25 male participants completed our experiments. 14 were flight nurses, and 11 were flight physicians. Their ages ranged from 24 to 56 years (Mean: 43.64; SD: 8.71).

### Environmental changes

The environmental conditions present on board the Dornier Do-328 Ambulance Jets were recorded for each test subject and waypoint. Means are displayed in table 2. Ambient cabin pressure was measured at mean 75.43 kPa (SD:1.38) when cruising altitude was reached and 75.94 kPa (SD:2.63) before descend, against normobaric conditions on ground levels (p< 0.001). To obtain a better understanding of these pressure values: 75 kPa correspond to an altitude of 2465 m above sea level. The subsequent reduction in partial oxygen pressure led to mild hypoxia in the test subjects. Mean oxygen saturations of the participants were measured at 92.92% (SD:2.00) after reaching cruising altitude and 93.6% (SD:1.93) before leaving cruising altitude – compared to a mean baseline saturation of 97.6% (SD:1.93, p< 0.001).

**Table 2.**
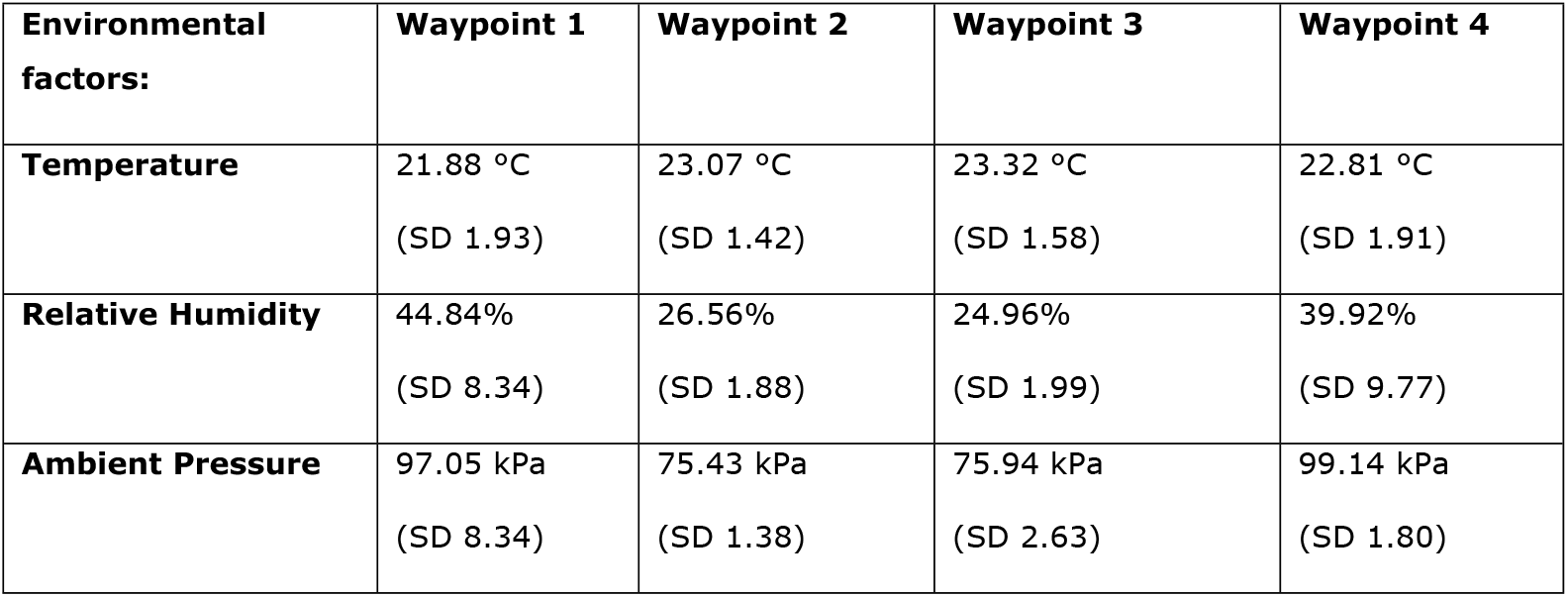
Environmental elements in effect during pain testing. Differences of statistical significance

### Somato-sensory testing – Nociception

The effects of environmental changes on perception and nociception are demonstrated by the Bland-Altman plots prepared for each test modality [12]. In this manuscript, we only display a short and exemplary collection of plots for the cold and heat pain thresholds (CPT and HPT) in Figures 2–5. Plots for all other analyzed modalities can be found as supplementary online content (Figures 6 - 23).

**Figure 2.**
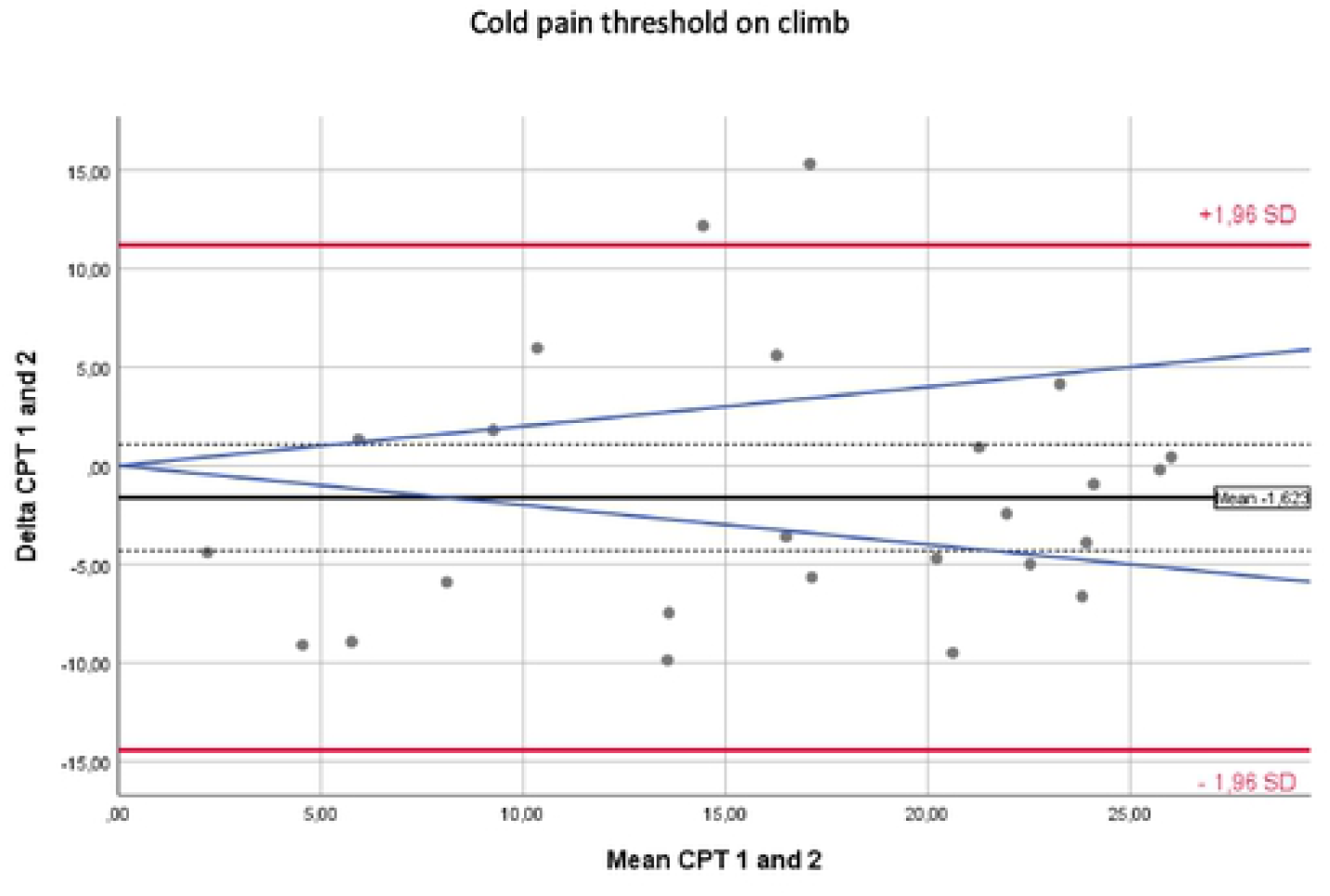
Bland Altman Plot - Cold Pain Threshold – Waypoints 1 against 2. The solid black lines in the Bland-Altman-Plots represent the mean of the differences. The confidence intervals for the means of differences are depicted as dashed black lines. The red upper (lower) lines show the upper (lower) limits of agreement equal to mean ± 1.96 SD. The blue lines represent a margin of ±20% around the means of measurements from each modality and serve as a possible indicator of clinical relevance.

**Figure 3.**
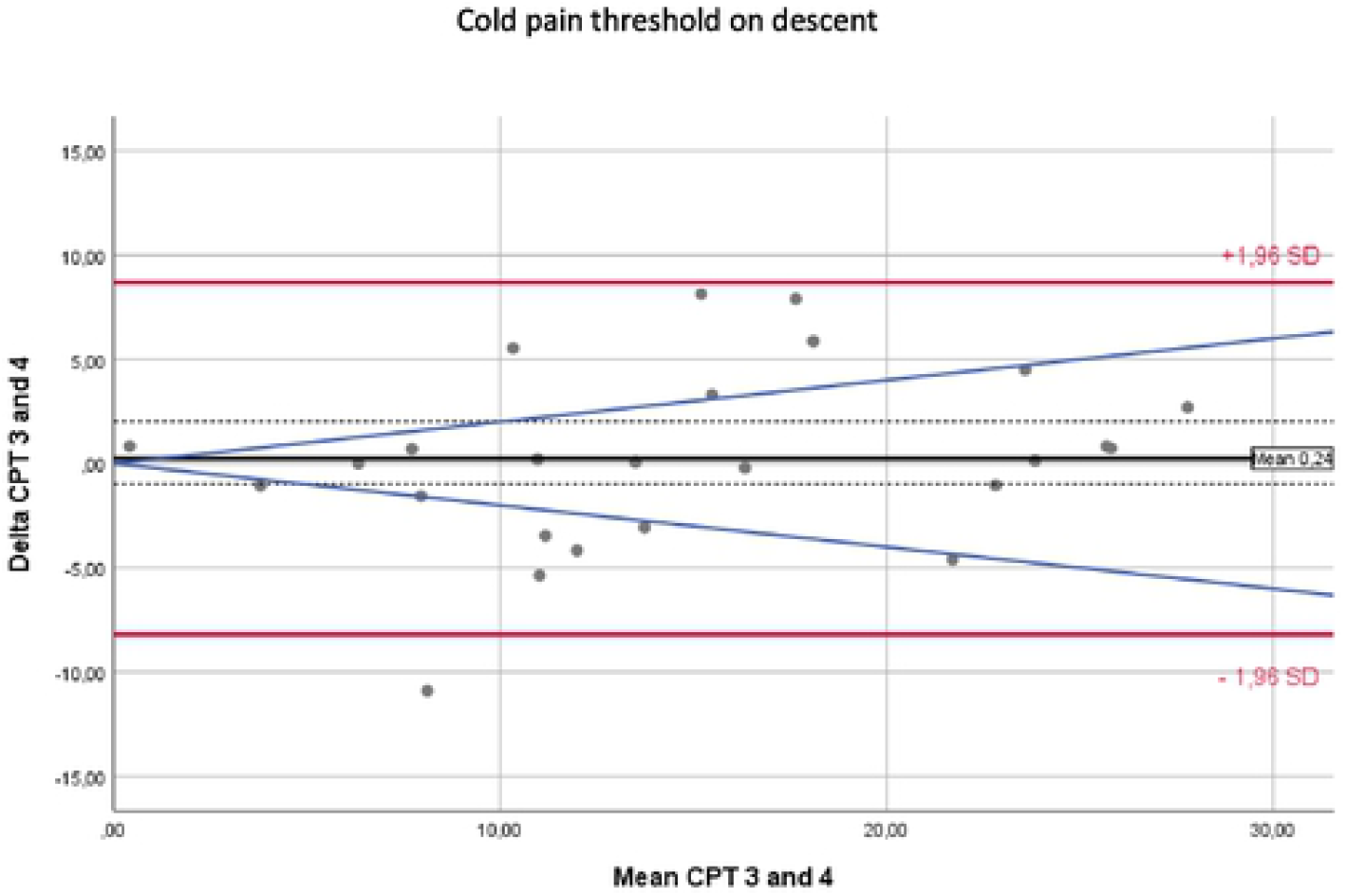
Bland Altman Plot - Cold Pain Threshold – Waypoints 3 against 4. The solid black lines in the Bland-Altman-Plots represent the mean of the differences. The confidence intervals for the means of differences are depicted as dashed black lines. The red upper (lower) lines show the upper (lower) limits of agreement equal to mean ± 1.96 SD. The blue lines represent a margin of ±20% around the means of measurements from each modality and serve as a possible indicator of clinical relevance.

**Figure 4.**
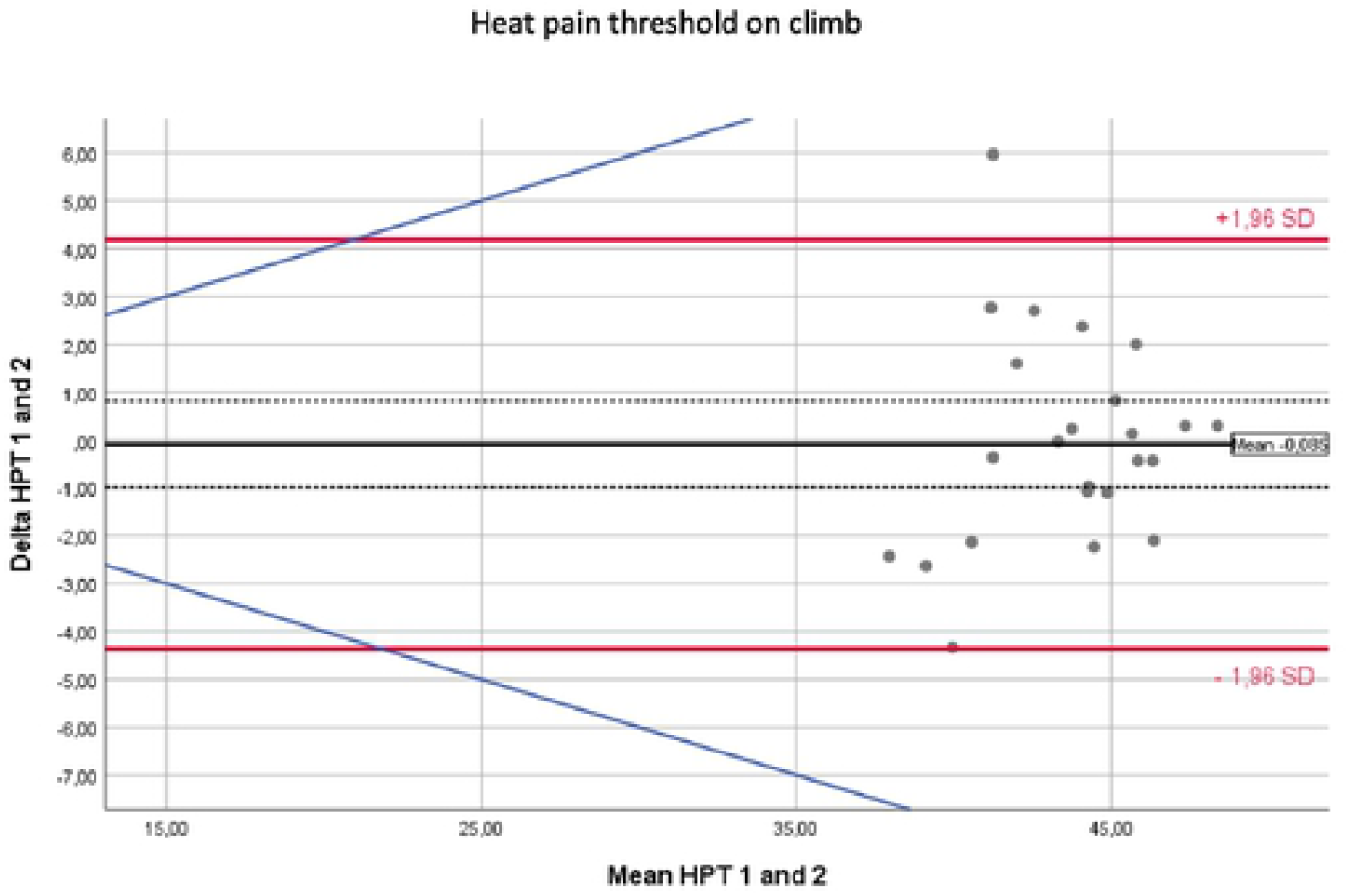
Bland Altman Plot - Heat Pain Threshold – Waypoints 1 against 2. The solid black lines in the Bland-Altman-Plots represent the mean of the differences. The confidence intervals for the means of differences are depicted as dashed black lines. The red upper (lower) lines show the upper (lower) limits of agreement equal to mean ± 1.96 SD. The blue lines represent a margin of ±20% around the means of measurements from each modality and serve as a possible indicator of clinical relevance.

**Figure 5.**
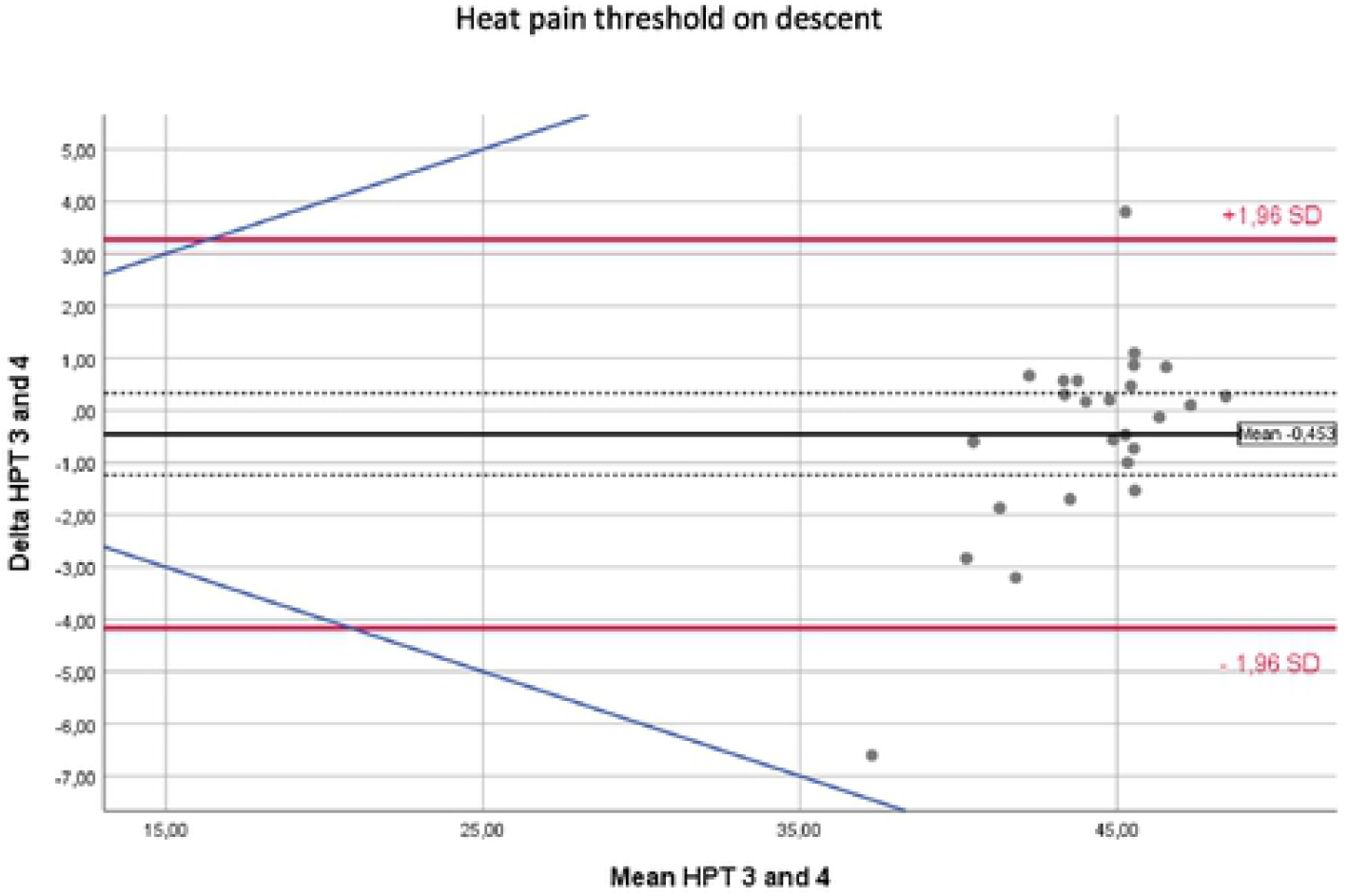
Bland Altman Plot - Heat Pain Threshold – Waypoints 3 against 4. The solid black lines in the Bland-Altman-Plots represent the mean of the differences. The confidence intervals for the means of differences are depicted as dashed black lines. The red upper (lower) lines show the upper (lower) limits of agreement equal to mean ± 1.96 SD. The blue lines represent a margin of ±20% around the means of measurements from each modality and serve as a possible indicator of clinical relevance.

Table 3 provides a comprehensive overview of each test and lists the means of the differences, their 95% confidence intervals and the limits of agreements (± 2SD) between measurements taken at waypoints 1 and 2. The estimated means of the differences were usually close to zero with some single differences demonstrating large nonsystematic fluctuations around this mean. This implies that environmental changes along the flight did not produce systematic bias but instead produced only random variations. Table 4 provides the same data for the comparison of waypoints 3 against 4.

**Table 3:** Means of differences, limits of agreement (LoA) and confidence intervals (CI) between pairs of measurements around changes in environmental conditions are shown for all analyzed sensory modalities. In this table between waypoints 1 and 2 (around take-off and climb).

| **Test modality** | **Unit** | **Mean of differences** | **CI mean** | **95% Upper LoA** | **CI upper limit of agreement** | **95% Lower LoA** | **CI lower limit of agreement** |
| --- | --- | --- | --- | --- | --- | --- | --- |
| **CDT12** | °C | -0.67 | -1.51 ; 0.16 | 3.29 | 1.61 ; 4.98 | -4.64 | -6.32 ; -2.95 |
| **HDT12** | °C | 0.16 | -0.56 ; 0.87 | 3.55 | 2.11 ; 5.00 | -3.24 | -4.68 ; -1.80 |
| **CPT12** | °C | -1.62 | -4.32 ; 1.07 | 11.18 | 5.74 ; 16.63 | -14.43 | -19.87 ; -8.99 |
| **HPT12** | °C | -0.09 | -0.98 ; 0.81 | 4.19 | 2.37 ; 6.00 | -4.36 | -6.17 ; -2.54 |
| **MPT12** | Nm | 16.51 | -0.47 ; 33.48 | 97.09 | 62.86 ; 131.33 | -64.08 | -98.31 ; -29.85 |
| **WUsS12** | NRS | 3.02 | -0.84 ; 6.89 | 21.38 | 13.58 ; 29.17 | -15.33 | -23.12 ; -7.53 |
| **WUmS12** | NRS | 1.01 | -1.96 ; 3.98 | 15.11 | 9.12 ; 21.10 | -13.09 | -19.08 ; -7.10 |
| **PPT12** | kg/cm<sup>2</sup> | 0.14 | -0.07 ; 0.34 | 1.12 | 0.70 ; 1.53 | -0.84 | -1.26 ; -0.43 |
| **PMDT12** | Intensity level | 0.40 | -0.22 ; 1.02 | 3.34 | 2.09 ; 4.59 | -2.54 | -3.79 ; -1.29 |
| **PMPT12** | Intensity level | 0.00 | -2.71 ; 2.71 | 12.89 | 7.41 ; 18.37 | -12.89 | -18.37 ; -7.41 |
| **PMAT12** | Intensity level | -0.98 | -4.36 ; 2.40 | 15.05 | 8.24 ; 21.86 | -10.20 | -23.82 ; -10.20 |

**Table 4:**
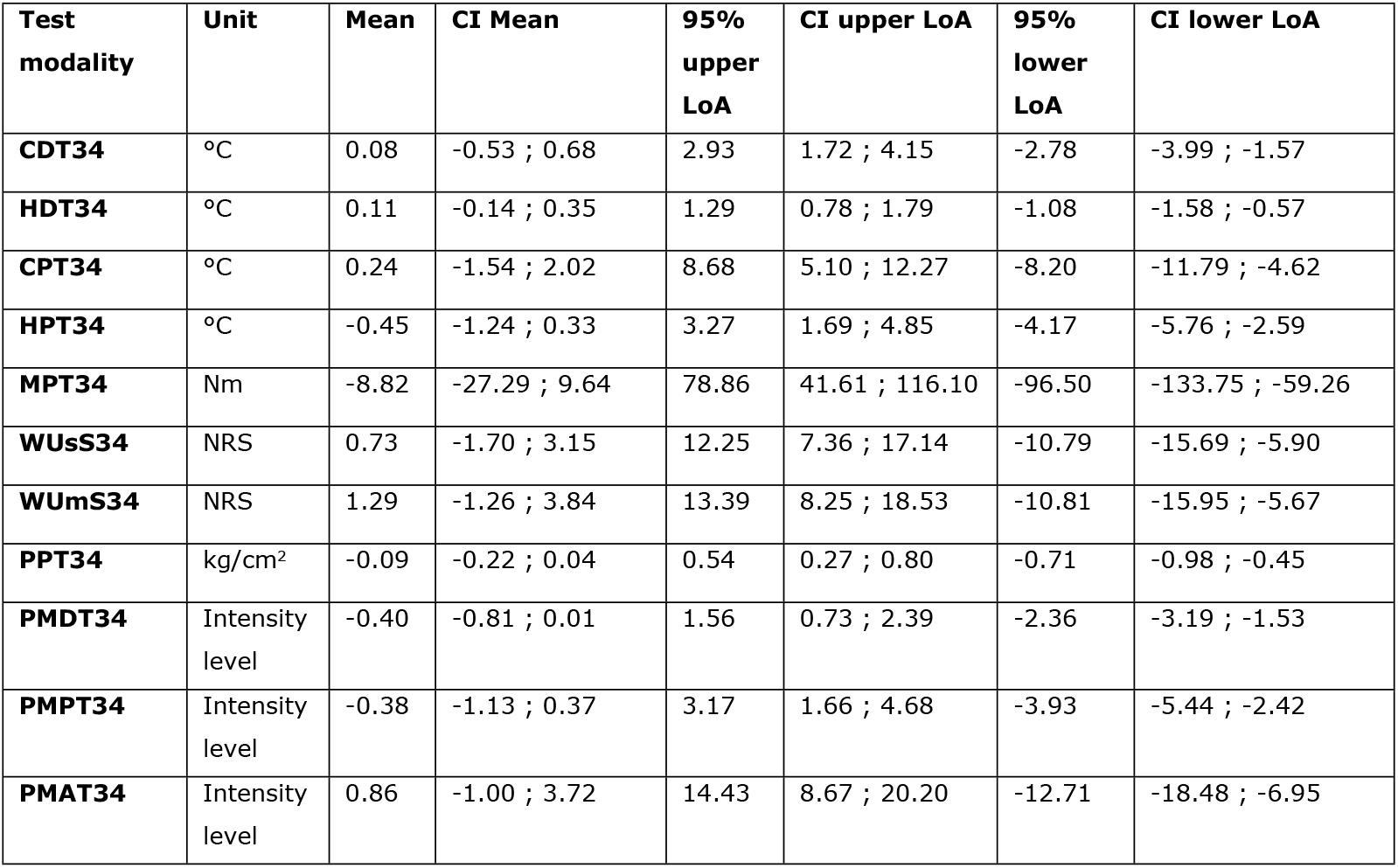
Means of differences, limits of agreement (LoA) and confidence intervals (CI) between pairs of measurements around changes in environmental conditions are shown for all analyzed sensory modalities. In this table between waypoints 3 and 4 (around descent and landing).

## Discussion

Long-distance inter-hospital transfers performed via fixed-wing air ambulances are frequent and steadily growing in number. Previous data from studies that investigated the impact of changing environmental conditions on neuro-sensory performance and nociception prompted us to suspect that patients undergoing transfers via airplanes could experience similar changes in nociception and that analgesia strategies may consequentially have to be re-evaluated. In this study, we present an elaborate test scenario aimed at assessing flight-related variations in perception and pain thresholds. Regarding the surrounding conditions, airplane travel is associated with large decreases in barometric pressure, partial oxygen pressure, and humidity as well as significant increases in vibration and noise exposure, all of which develop over very short time spans. Our data suggest that despite these significant and systematic environmental changes, the variations in nociception that occur during an ambulance flight are nonsystematic and random – according to our comprehensive scope of sensory modalities. Variations in the detected differences against their means could occur in a larger extent in a number of cases. (I.e., measurements lying outside the blue 20% margin in each plot.) In some of the tested modalities, more than half of the test subjects displayed means of differences exceeding 20%, which is a – certainly debatable – margin of clinical significance. However, these cases occurred without any clear pattern, and no systematic routine allowed us to predict the direction of an individual’s change in nociception, or whether he might not be affected at all by the environmental stressor he was exposed to.

While our study is not a final assessment that should be used to guide analgesia in a systematic way (e.g. more or less dosing), we conclude that flight-related changes in the environment have the potential to erratically influence some individuals’ nociception. This finding calls for increased clinical suspicion of altered, whether higher or lower - analgesia requirements during those phases of a transfer we tested, during which external conditions change profoundly. Our data indicate that repeated pain assessments should potentially be carried out at times such as take-off and landing in patients requiring analgesia.

### Findings in the context of previous data

As described in the introduction section, previous studies have presented data that suggest that the environment can have systemic effects on nociception. At first glance, our findings seem to contradict these studies, but a closer look allows a reconciliation of their conclusions and ours. Regarding the effects of high altitude on mountaineers, it must be acknowledged that the environmental conditions experienced on board airplanes are not as extreme as those experienced in the Himalayans and that the duration of exposure was considerably shorter for our volunteers [4]. In fact, the experiments investigating the aggravating effects of short-term weather changes on neuropathic pain in guinea pigs correspond somewhat more closely to our setting of environmental changes [3]. However, a fundamental difference between our test setting and the one used with the guinea pigs is the pathophysiological condition of the test subjects. In contrast to the test animals, which suffered from neuropathic pain, our volunteers were healthy individuals without any pain other than that caused by the mild stimuli of the QST battery. It is conceivable, that in order to be influenced by external environmental stressors such as those used in our study, pain must be present as an actual and persistent disorder, not just as a brief experimental stimulus. In the end, it seems worth considering whether the effects of environmental changes on a cohort of test subjects who actually experiencing pain should be investigated before we reject the hypothesis that flight-related environmental changes have relevant effects on human nociception. After all, pain can, in itself, have systemic effects on stimulus detection and pain thresholds and can lead to very complex but distinct secondary disorders, such as hyperalgesia and allodynia [13–15].

### Strengths

In our study we conducted nociception testing in a real-life in-flight setting unparalleled by that used in any previous study. All external factors present during the actual transfers of patients were also present under our experimental conditions. This included factors that are easy to simulate and easy to measure such as barometric pressure as well as factors that are more difficult to replicate under laboratory conditions, such as motion, vibrations, noise, odors, and others - some of which we might not even be aware of as to their existence.

We demonstrate the feasibility of using a complex and comprehensive somato-sensory assessment in a unique surrounding area. Our study clears the way for further investigations of nociception in selected, clinically relevant subpopulations submitted to flight conditions. As mentioned above, the fact that our findings do not support the notion of a systematic effect on nociception in healthy volunteers does not exclude the possibility that such effects could occur in individuals actually experiencing pain at the time of transfer.

### Limitations

Naturally, these strengths stand vis-à-vis with several limitations. First, the small sample-size of our study population is the most relevant limitation. Second, due to the make-up of the work force of the ADAC Air Ambulance, our study population consisted only of men in young adulthood or middle age. This selection bias limits the generalizability of our findings as age and sex are factors that are known to influence nociception [16–18]. Selection may also have been affected by the so-called healthy worker effect [19]. It is conceivable that those individuals who are actually affected the most by flight-related environmental changes would not work in the field of aeromedical retrievals and that we therefor inadvertently tested a subpopulation of (in a manner of speaking) immune individuals.

## Conclusions

Air ambulance flights submit patients to extraordinary and rapidly changing environmental conditions, and providers of care and researchers have aimed to explore the effects of airplane travel on patients. In this study, we investigated the feasibility of somatosensory testing on the basis of QST to identify possible flight-related changes in stimulus perception and pain thresholds. In consideration of the declared limitations, we can present several novel findings. We demonstrate the feasibility of using a complex and comprehensive method of nociceptive testing under real-life in-flight conditions. This opens up the possibility that future investigations could explore nociception among patients who require analgesia, for whom we must strive to optimize our provision of care. However, with regard for our healthy volunteers, perception thresholds, pain thresholds, and above-threshold pain were not subject to systematic effects along the major changes of the environment accompanying the different stages of an ambulance flight. Nociception was considerably altered in a relevant percentage of individuals, but our data do not suggest a methodical way to predict such occurrences.

## Supporting information

Bland-Altman Plots for tested somato-sensory modalities other than cold and heat pain thresholds are available as supplemental online content.

## Abbreviations

ADAC: Allgemeiner Deutscher Automobil Club
CDT: Cold Detection Threshold
CI: Confidence Interval
CPT: Cold Pain Threshold
DOI: Digital Object Identifier
HDT: Heat Detection Threshold
HPT: Heat Pain Threshold
ICU: Intensive Care Unit
LoA: Limit of Agreement
MPT: Mechanical Pain Threshold
NRS: Numerical Rating Scale
PMAT: Pain Matcher Abort Threshold
PMDT: Pain Matcher Detection Threshold
PMPT: Pain Matcher Pain Threshold
PPT: Pressure Pain Threshold
QST: Quantitative Sensory Testing
SD: Standard Deviation
WumS: Wind Up Multiple Stimuli
WusS: Wind Up Single Stimulus

## Declarations

### Ethics approval and consent to participate

This study was evaluated and approved by the University of Erlangen-Nuremberg’s ethics council beforehand. (Decision number 81_13 B.) Informed written consent was obtained from each participant before testing.

### Consent for publication

All authors read and approved the final version of the manuscript.

### Availability of data and material

The datasets generated and analyzed during the current study are available in the Zenodo data repository: 10.5281/zenodo.2563176

### Authors’ contributions

JP conceived of and co-conducted the study, co-performed the statistical analysis and is the main author of the manuscript. SE co-conducted the study, co-performed the statistical analysis and revised the manuscript critically for its intellectual content. AW, AM, JS and MM helped in the execution of the study and revised the manuscript critically for its intellectual content.

## Acknowledgments

We thank Clemens Forster, Gabriele Göhring-Waldeck and Roland Schiffmann for their excellent assistance in the logistical preparations of our study. Finally, we would like to thank all of the volunteers for their contributions to our study.

